# CD4^+^ T cells recruit, then engage macrophages in cognate interactions to clear *Mycobacterium tuberculosis* from the lungs

**DOI:** 10.1101/2024.08.22.609198

**Authors:** Samuel H Becker, Christine E Ronayne, Tyler D Bold, Marc K Jenkins

## Abstract

IFN-γ-producing CD4^+^ T cells are required for protection against lethal *Mycobacterium tuberculosis* (*Mtb*) infections. However, the ability of CD4^+^ T cells to suppress *Mtb* growth cannot be fully explained by IFN-γ or other known T cell products. In this study, we show that CD4^+^ T cell-derived IFN-γ promoted the recruitment of monocyte-derived macrophages (MDMs) to the lungs of *Mtb*-infected mice. Although the recruited MDMs became quickly and preferentially infected with *Mtb*, CD4^+^ T cells rapidly disinfected the MDMs. Clearance of *Mtb* from MDMs was not explained by IFN-γ, but rather by MHCII-mediated cognate interactions with CD4^+^ T cells. These interactions promoted MDM expression of glycolysis genes essential for *Mtb* control. Thus, by recruiting MDMs, CD4^+^ T cells initiate a cycle of bacterial phagocytosis, *Mtb* antigen presentation and disinfection in an attempt to clear the bacteria from the lungs.

## Introduction

*Mycobacterium tuberculosis* (*Mtb*) is a phagosomal pathogen that infects the lungs and is responsible for tuberculosis. While natural sterilizing immunity to *Mtb* is rare, the majority of immunocompetent individuals generate sufficient immunity to prevent disease [1]. This immunity depends on CD4^+^ T cells as evidenced by the fact that people and primates with CD4^+^ T cell deficiency due to immunodeficiency virus infection, and mice genetically deficient or depleted of CD4^+^ T cells, are hyper-susceptible to tuberculosis [2–6].

During infection, naïve CD4^+^ T cells that express T cell antigen receptors (TCRs) specific for major histocompatibility complex class II (MHCII)-bound peptides from the infecting microbe become activated in secondary lymphoid tissue and proliferate to form a large population of effector cells. These effector cells then differentiate based on cytokines made by cells of the innate immune system. Pathogens that infect the phagosomes of myeloid cells stimulate the infected cells and other innate immune cells to produce IL-12, which instructs pathogen-specific effector cells to differentiate into T helper type 1 (Th1) cells that are defined by the capacity to make interferon gamma (IFN-γ) [7]. The Th1 cells subsequently traffic to infected tissues to promote clearance of the infection.

The mechanism, however, that CD4^+^ T cells use to control *Mtb* infection in the lungs is unclear. Humans with genetic deficiencies in IL-12 or IFN-γ signaling succumb to lethal infections by environmental mycobacteria as well as *Mycobacterium bovis* BCG, an attenuated strain that has been used as a *Mtb* vaccine. Surprisingly, however, there is a weaker correlation between IL-12 or IFN-γ signaling defects and lethal *Mtb* infections [2, 8]. In addition, although mice lacking an intact IFN-γ receptor, as well as mice whose CD4^+^ T cells cannot make IFN-γ, die from infection more quickly than wild-type (WT) mice, the ability of CD4^+^ T cells to directly restrict *Mtb* growth within the lungs is primarily independent of IFN-γ production [9–13]. In addition, other Th1 products TNF-α, perforin, FAS, and GM-CSF fail to explain CD4^+^ T cell-mediated control of *Mtb* replication *in vivo* [9, 11, 13]. Thus, although there is no doubt that CD4^+^ T cells are critical for control of *Mtb* infection, the mechanism of action has not been resolved.

Regardless of the mechanism, it is likely that CD4^+^ T cells clear *Mtb* by inducing an antimicrobial state in the cells that contain bacteria. *Mtb* primarily resides in the lungs within neutrophils (PMNs) and a population of poorly characterized cells with a macrophage or dendritic cell (DC)-like phenotype that are commonly referred to as interstitial macrophages (IMs) [14–18]. IMs express high levels of MHCII and thus would be capable of presenting MHCII-bound *Mtb* peptides to lung CD4^+^ T cells after engulfing *Mtb*.

Experiments with mixed bone marrow chimeric mice demonstrated that MHCII-deficient IMs contain more *Mtb* bacilli than WT IMs within the same animal [19]. These observations suggest that CD4^+^ T cell recognition of *Mtb* peptide:MHCII complexes presented by infected IMs enhances the capacity of the IMs to kill the bacteria within them. The nature of this enhanced capacity is unknown.

In this study, we definitively show that IMs within *Mtb*-infected lungs are of monocyte origin, and thus refer to them as monocyte-derived macrophages (MDMs). We determined that MDM recruitment to *Mtb*-infected lungs is primarily not an innate immune response, but rather is driven by IFN-γ from CD4^+^ T cells. Once recruited, MDMs became infected at a high rate but underwent disinfection that was dependent on expression of MHCII. Antigen presentation was required for MDMs to fully upregulate glycolysis genes known to be critical for the suppression of intracellular *Mtb* growth. Thus, Th1 cells respond to intracellular bacteria by recruiting phagocytic MDMs that act as antigen-specific targets for Th1 effector function.

## Results

### Identification of monocyte-derived macrophages (MDMs) in *Mtb*-infected mouse lungs using CX3CR1 and Ly-6C

The myeloid cells that were infected in the lungs of *Mtb*-infected mice were studied with a goal of clarifying the identity of IMs. C57BL/6 mice were challenged with a low dose of aerosolized *Mtb* (∼100 bacilli/mouse) and the lungs were analyzed at 3 weeks post-infection using flow cytometry. After excluding T and B cells, canonical markers were used to identify PMNs (Ly-6G^+^ CD11b^+^), alveolar macrophages (SIGLEC-F^+^ CD11c^+^), and NK cells (NK1.1^+^) (**Fig. 1A**). Among the remaining cells, CX3CR1 was used as a marker to distinguish monocytes from macrophages and DCs. A large population of CX3CR1^hi^ cells that contained two subsets was detected: Ly-6C^hi^ CD11c^—^ cells, provisionally identified as classical monocytes (CMs), and Ly-6C^lo/int^ CD11c^lo/int^ cells, provisionally identified as nonclassical monocytes (NCMs). CX3CR1^lo^ cells could also be divided into two subsets: Ly-6C^hi^ CD26^lo^ cells, provisionally identified as monocyte-derived macrophages (MDMs), and Ly-6C^lo^ CD26^hi^ cells, provisionally identified as DCs (**Fig. 1B**).

**Figure 1.**
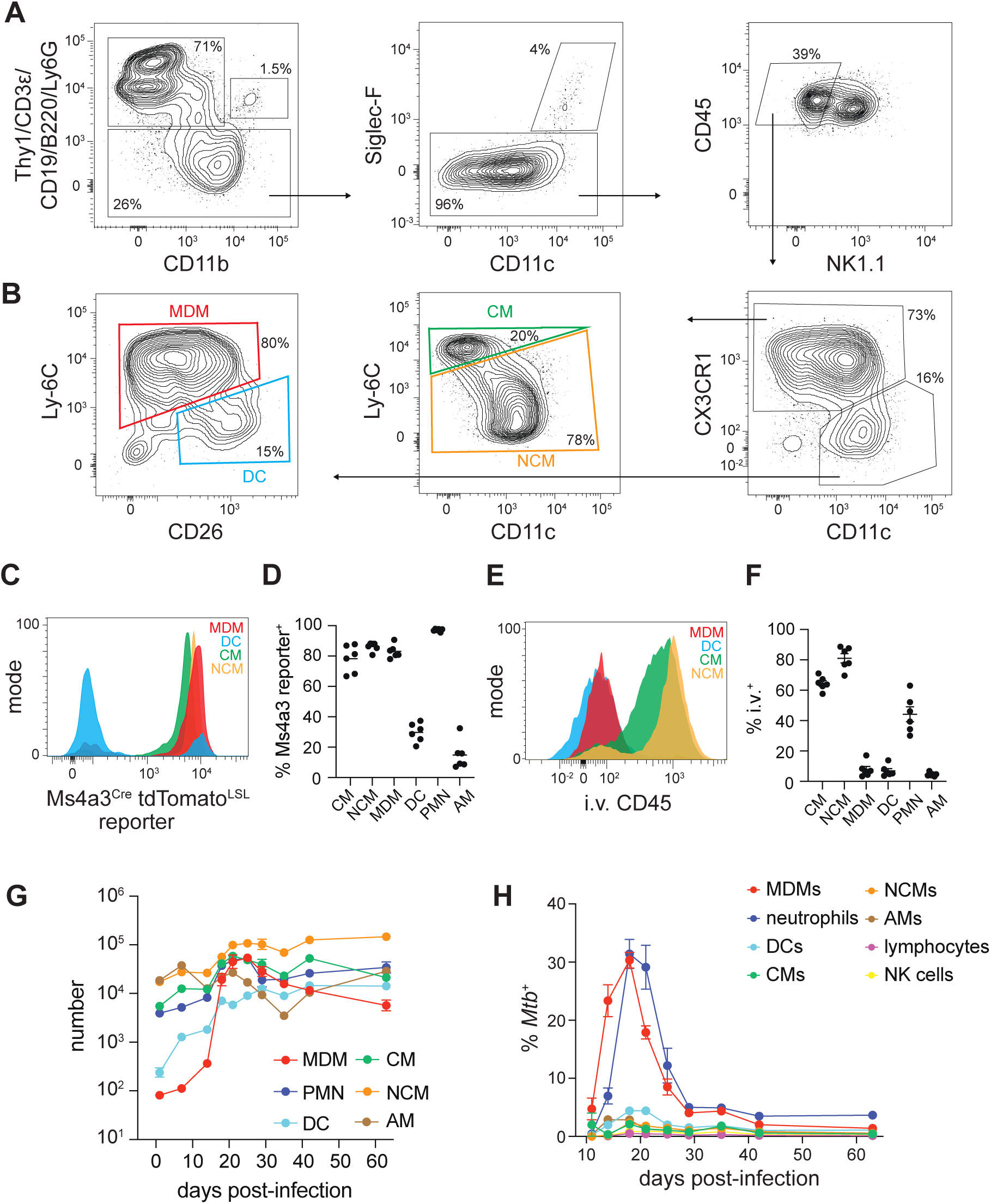
CX3CR1^lo^ CD11c^hi^ CD26^lo^ Ly-6C^hi^ monocyte-derived macrophages (MDMs) are recruited, preferentially infected, and disinfected. (A) and (B) Cells in the lungs of WT mice at 3 weeks post-infection with *Mtb*, pre-gated on CD45^+^ singlets. MDM, monocyte-derived macrophage; DC, dendritic cell; CM, classical monocyte; NCM, nonclassical monocyte. (C-D) tdTomato fluorescence among indicated cell populations. PMN, neutrophil; AM, alveolar macrophage. (E-F) Staining of indicated lung populations with intravascular (i.v.) CD45 antibody. (G) Number of indicated lung populations by subset. (H) Frequency of *Mtb*-mScarlet-infected cells within each of the indicated lung populations. Data in (A) through (F) were collected at 3 weeks post-infection.

We next performed tests to validate the identity of each population. NCMs, which differentiate from CMs, downregulate Ly-6C and upregulate another surface protein, Treml4 [20]. CX3CR1^hi^ Ly-6C^lo/int^ CD11c^lo/int^ cells, putatively identified as NCMs, expressed uniformly high levels of Treml4 compared to the remaining cells (**Fig. S1A**). NCMs are also acutely dependent on the growth factor CSF-1 for survival [21]. When *Mtb* infected mice were treated for 1 week with an antibody that blocks CSF-1 signaling (α-CD115), the NCMs were preferentially depleted compared to CMs and MDMs (**Fig. S1B**). Together, the results indicated that CX3CR1^hi^ Ly-6C^lo/int^ CD11c^lo/int^ cells were NCMs.

Previous studies indicate that IMs and DCs within the lungs both express CD64 and MHCII, making these cells difficult to resolve [22]. In line with those observations, the cells provisionally identified as MDMs and DCs expressed similarly high levels of CD64 and MHCII compared to CMs and NCMs (**Fig. S1C**). However, the CX3CR1^lo^ population could be separated into Ly-6C^lo^ CD26^hi^ DCs and Ly-6C^hi^ CD26^lo^ MDMs (**Fig. 1B**). The identity of the MDMs and DCs was further tested using lineage-tracing reporter mice. *Ms4a3* is expressed by cells of the monocyte and granulocyte lineages, but not in classical DCs (cDCs) or non-hematopoietic macrophages such as AMs [23]. Ms4a3^Cre^ tdTomato^LSL^ mice, in which cells of monocyte or granulocyte origin express tdTomato, were infected with *Mtb* and assayed at 3 weeks post-infection to assess the origins of the lung myeloid populations. tdTomato was uniformly expressed by MDMs, CMs and NCMs, but not DCs (**Fig. 1C**). Thus, the cells that were identified as MDMs indeed arise from the monocyte lineage.

Only about 30% of DCs were tdTomato^+^ indicating that a small number of DCs are monocyte-derived or the *Ms4a3* reporter is expressed in an off-target fashion in some DCs. As a control for the fidelity of reporter expression, most AMs were tdTomato^—^, reflecting their non-hematopoietic origin, while all PMNs were tdTomato^+^ (**Fig. 1D**).

Location in blood was also assessed to validate the cell identifications. Intravenous (i.v.) injection of a CD45 antibody labeled most of the cells that were identified as CMs and NCMs, consistent with these cells being blood monocytes. By contrast, nearly all MDMs and DCs were not labeled with i.v. CD45 antibody, indicating tissue residence (**Fig. 1E**). Many PMNs were also labeled with i.v. CD45 antibody while AMs were not, consistent with their location in blood and tissue, respectively (**Fig. 1F**). Overall, the i.v. labeling and phenotyping results demonstrate CX3CR1, CD11c, Ly-6C and CD26 effectively resolve CMs, NCMs, MDMs and DCs in murine lungs. Importantly, MDMs had high CD64 and MHCII expression previously attributed to IMs.

### MDMs are recruited to the lungs and undergo rapid infection followed by disinfection

The number of myeloid cells in the lungs were then measured after *Mtb* infection. CMs, NCMs, AMs, and PMNs accounted for most of the myeloid cells in the lungs of uninfected mice. After infection, the cells in these populations increased slowly if at all and no more than 10-fold over a 2 month period. By contrast, very few DCs or MDMs were detected before infection and both populations increased greatly after infection. The increase in lung MDMs was especially dramatic. The number of MDMs slowly increased between days 0 and 14, then increased sharply and peaked on day 21 after a 100-fold increase. The MDMs then remained elevated through day 63 (**Fig. 1G**). Thus, events occurring between 14 and 21 days post-infection drove the recruitment of most of the MDMs to the lungs.

The frequency with which each population was infected with *Mtb* was also measured using an mScarlet-expressing *Mtb* strain. MDMs and PMNs were much more likely to contain intracellular *Mtb* than the other myeloid cell populations. At 14 days post-infection, MDMs became heavily infected while PMNs remained mostly uninfected. On day 18, the infectious burden of both populations peaked at approximately 30 percent. On day 21, MDM infection had sharply decreased while PMN infection remained at 30 percent. By day 28, less than 5 percent MDMs and PMNs were infected, and this low infectious burden was stable up to day 63 (**Fig. 1H**). Overall, our results indicated that MDMs are recruited during the third week of infection, become rapidly infected, and undergo similarly rapid disinfection based on the frequency of mScarlet^+^ MDMs. By contrast, PMNs are relatively abundant from the onset of infection, become infected slightly later than MDMs, and are disinfected more slowly than MDMs.

### CD4^+^ T cells promote both MDM recruitment and MDM disinfection

Work by others showing that *Mtb*-specific CD4^+^ T cells migrate into the lungs between 14 and 21 days post-infection [24] raised the possibility that these lymphocytes are responsible for MDM recruitment. This hypothesis was tested by using fluorescently-labeled I-A^b^ (the MHCII molecule of C57BL/6 mice) tetramers containing a peptide from the *Mtb* protein ESAT6 (ESAT6p:I-A^b^) and flow cytometry to track the emergence of *Mtb-*specific CD4^+^ T cells in the lungs [25] (**Fig. S2A**) over time after infection. Although *Mtb* CFUs increased logarithmically starting from the onset of infection, the abundance of ESAT6p:I-A^b^-specific CD4^+^ T cells and MDMs was very low until day 14 and then began increasing at a high rate thereafter (**Fig. 2A**). These results are consistent with the hypothesis that MDM recruitment to the lungs is primarily mediated by *Mtb*-specific CD4^+^ T cells rather than earlier innate immunity.

**Figure 2.**
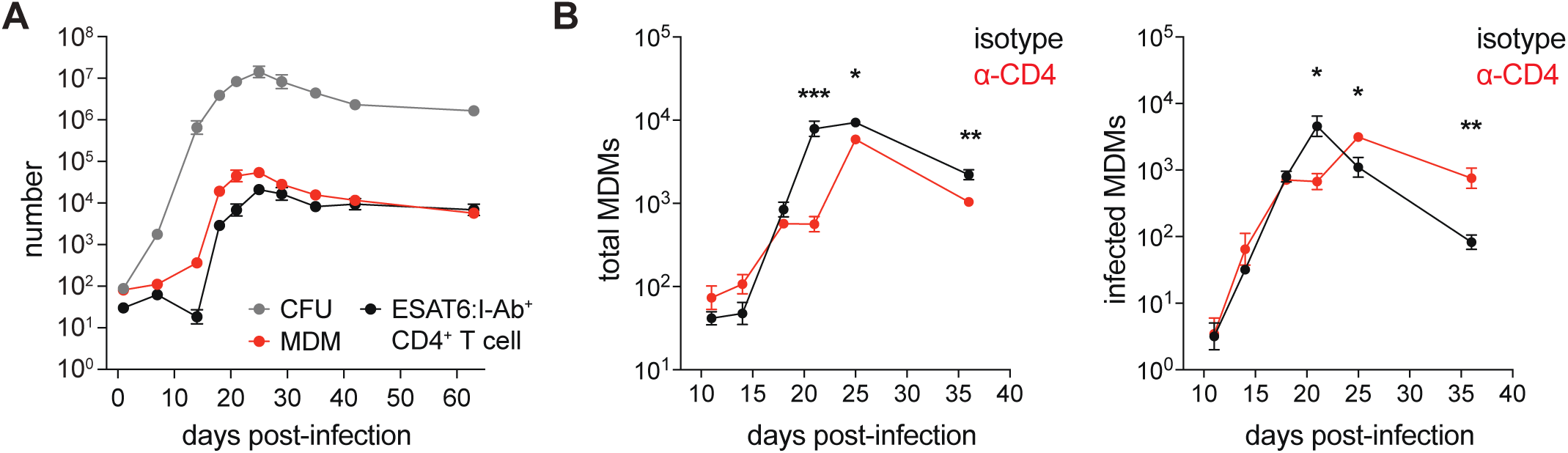
CD4^+^ T cells promote early MDM recruitment and MDM disinfection. (A) Number of *Mtb* colony-forming units (CFUs), MDMs, and CD4^+^ T cells staining positively for ESAT6:I-A^b^ tetramer in the lungs. (B) Total MDMs in the lungs of mice treated CD4^+^ T cell-depleting (α-CD4) or isotype control antibodies. (C) Number of infected MDMs in the lungs of mice treated with α-CD4 or isotype control antibodies.

The role of CD4^+^ T cells in the recruitment of MDMs to the lung was tested by treating mice with an antibody (α-CD4) that completely depletes CD4^+^ T cells (**Fig. S2B**). CD4^+^ T cell sufficient mice treated with isotype control antibody experienced a sharp increase in total MDMs starting on day 14 that plateaued on day 21 (**Fig. 2B**) as observed in untreated mice **(Fig. 1G)**. *Mtb*-infected MDMs also increased in CD4^+^ T cell sufficient mice until day 21, but then declined 50-fold by day 36 (**Fig. 2C**). By contrast, the lungs of CD4^+^ T cell-depleted mice had approximately 10-fold fewer total and *Mtb*-infected MDMs on day 21 than control mice (**Figs. 2B-C**). Thus, the recruitment of MDMs in control mice around day 21 was CD4^+^ T cell-dependent and the recruited MDMs became infected. The number of *Mtb*-infected MDMs in CD4^+^ T cell-depleted mice, however, was similar between days 21 and 36, a period when the number decreased 50-fold in control mice (**Fig. 2C**). These results indicate that CD4^+^ T cells perform two temporally distinct functions: initial recruitment of infection-prone MDMs into the lungs and then disinfection of the recruited MDMs that become infected.

### CD4^+^ T cell-derived IFN-γ causes MDM recruitment

The mechanism of the early CD4^+^ T cell-dependent recruitment of MDMs into the lungs was then explored. CMs, which are generated in the bone marrow before entering the bloodstream, are the precursors to lung-resident MDMs [26, 27]. It was possible that CD4^+^ T cell depletion diminished the number of CMs in the blood that could be recruited into the lungs. However, although a-CD4 treatment reduced the number of MDMs (**Fig. 3A**), which the i.v. CD45 antibody experiments showed were in the lung tissue **(Fig. 1E**), it had much less effect on the CMs (**Fig. 3A**) that were in the blood in the lung samples (**Fig. 1E**). It was therefore possible that CD4^+^ T cells cause CMs to migrate from the blood into the lung tissue and differentiate into MDMs. This hypothesis was tested by monitoring the fate of purified CMs (**Fig. S3A**) 18 hours after adoptive transfer into day 20 *Mtb*-infected mice depleted of CD4^+^ T cells (**Fig. 3B**). The a-CD4-treated mice contained 10-fold fewer donor CM-derived MDMs than isotype control antibody-treated mice (**Fig. 3C**), demonstrating that CD4^+^ T cells were required for blood CMs to become lung MDMs.

**Figure 3.**
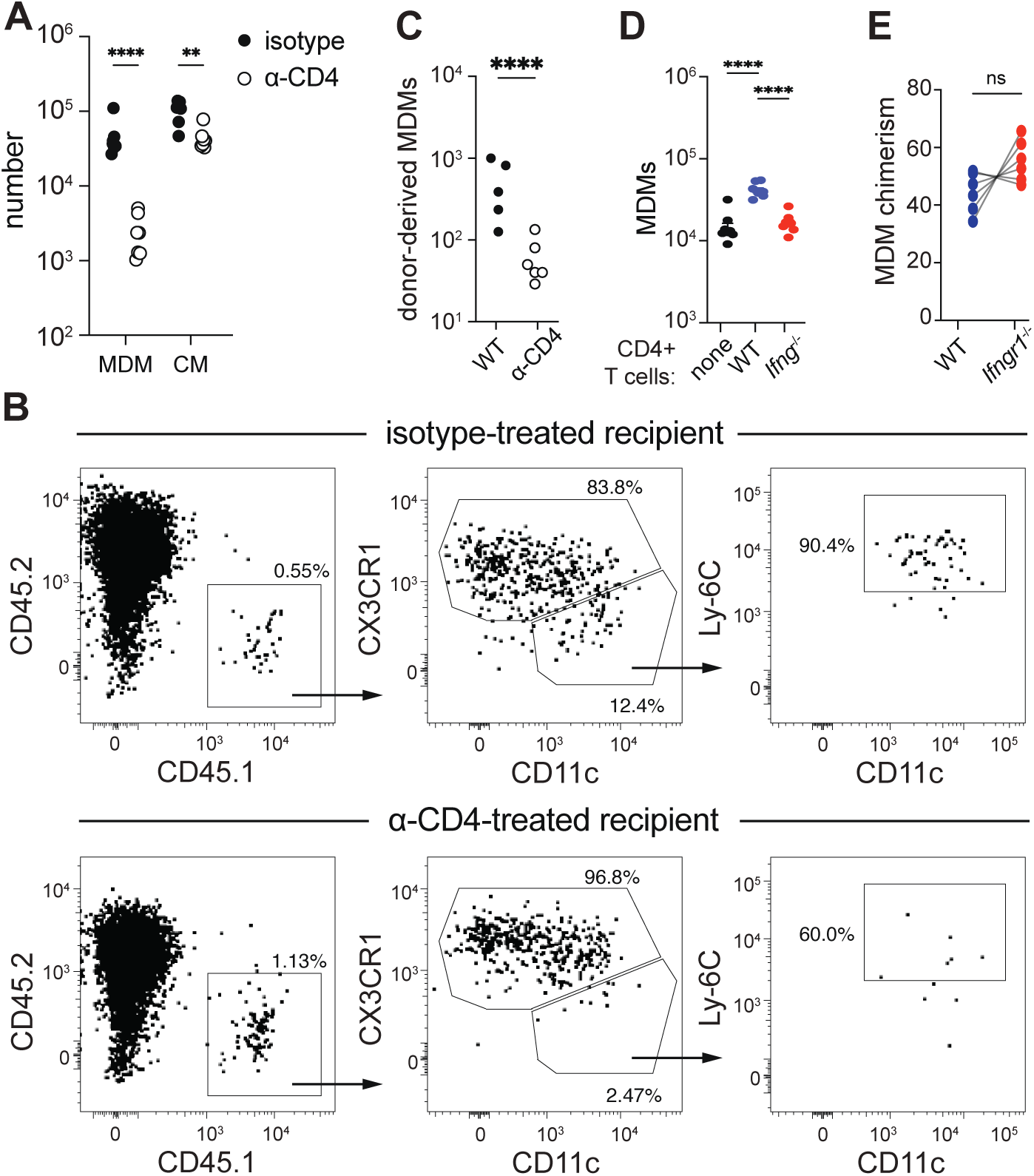
CD4^+^ T cell-derived IFN-γ recruits lung MDMs from blood CMs. (A) Number of indicated cells in the lungs of α-CD4- or isotype control-treated mice at 3 weeks post-infection. (B) Phenotype of CMs purified from naïve CD45.1^+^ donor mice 24 hours after i.v. injection into day 20 *Mtb*-infected CD45.2 recipient mice. (C) Number of donor CM-derived MDMs in the lungs of mice from (B). (D) Number of MDMs in the lungs of T cell-deficient mice given no CD4^+^ T cells, WT CD4^+^ T cells, or IFN-γ-deficient (*Ifng*^—/—^) CD4^+^ T cells and infected with *Mtb* for 4 weeks. (E) Frequency of WT versus IFN-γ receptor-deficient (*Ifngr1*^—/—^) MDMs in the lungs of mice with a 1:1 mix of WT and IFN-γ receptor-deficient bone marrow at 4 weeks post-infection. Lines indicate paired data for each mouse.

The properties of CD4^+^ T cells that promote the recruitment of MDMs were then assessed. IFN-γ production was a prime candidate as this is a major function of Th1 cells. Mice were generated with or without the *Ifng* gene in CD4^+^ T cells by transferring CD4^+^ T cells from naïve WT or *Ifng*^—*/*—^ donor mice into T cell-deficient (*Tcra*^—*/*—^) recipients. Both groups of CD4^+^ T cell transfer recipients, as well as a control group of T cell-deficient mice receiving no transferred cells, were then infected with *Mtb* to assess IFN-γ levels and MDM abundance in the lungs. At 4 weeks post-infection, T cell-deficient mice that received WT donor CD4^+^ T cells contained significantly more lung IFN-γ than mice that received IFN-γ-deficient CD4^+^ T cells or no CD4^+^ T cells (**Fig. S3B**). Furthermore, T cell-deficient mice that received WT CD4^+^ T cells contained significantly more lung MDMs than T cell-deficient mice that received no CD4^+^ T cells, and this increase in MDMs was completely abolished in T cell-deficient mice that received IFN-γ-deficient CD4^+^ T cells (**Fig. 3D**). These results indicate that CD4^+^ T cells recruit MDMs by secreting IFN-γ. The eventual recovery of MDMs in a-CD4-treated mice (**Fig. 2B**) is likely explained by IFN-γ-producing CD8^+^ T cells, which are known to arise in CD4^+^ T cell-deficient mice approximately 1 month after infection [5].

IFN-γ has not been previously shown to have chemotactic properties or to induce the differentiation of CMs into MDMs. To determine if direct IFN-γ signaling to blood CMs was required for their migration into the lungs or their differentiation into MDMs, mixed bone marrow chimeric mice were generated by reconstituting irradiated mice with a 1:1 ratio of WT (CD45.1^+^) and *Ifngr1*^—*/*—^ (CD45.2^+^) bone marrow cells. The chimeric mice were infected with *Mtb* and MDMs were identified in the lungs at 3 weeks post-infection (**Fig. S3C**). Not only were the MDMs derived from the IFNgR1-deficient bone marrow cells not reduced, but they were greater in number than the MDMs derived from WT bone marrow cells (**Fig. 3E**), perhaps due to the loss of the anti-proliferative effect of IFN-γ [28]. Thus, although CD4^+^ T cell-derived IFN-γ promoted the accumulation of MDMs in the lungs, expression of the IFN-γ receptor by the MDMs or their precursors was not. Instead, CD4^+^ T cell-derived IFN-γ must promote MDM recruitment indirectly by signaling to other cells.

### Antigen presentation by CCR2-dependent cells is required for CD4^+^ T cell-mediated immunity to *Mtb*

The mechanism by which CD4^+^ T cells disinfect the recruited MDMs was then addressed. MDMs are poised for this purpose because of the ability to present MHCII-bound *Mtb* peptides and thus directly receive activating signals from cognate CD4^+^ T cells. *CCR2^CreERT2-GFP/WT^ H2-Ab1^flox/flox^* (MHCII^ΔCCR2^) mice, in which tamoxifen (TAM) treatment causes cells expressing CCR2 to delete the gene encoding the I-A^b^ beta chain and express GFP [29], were used to address this possibility. TAM treatment is expected to result in loss of MHCII expression in MDMs in these mice because CCR2 is expressed in the CM precursors of MDMs.

Importantly, prior reports suggest that antigen presentation by CCR2-expressing cells is not required for the initial priming of *Mtb*-specific CD4^+^ T cells in lymph nodes [30] and thus this approach should eliminate antigen presentation by CCR2^+^ cells in the lungs while retaining *Mtb*-specific CD4^+^ T cell activation in lymph nodes.

MHCII^ΔCCR2^ and Cre^−^ littermates (MHCII^WT^) were infected with *Mtb* and given TAM-containing chow starting on the day of infection. The MDMs of MHCII^ΔCCR2^ mice lacked MHCII 3 weeks post-infection and expressed GFP as expected. The DCs of MHCII^ΔCCR2^ mice also lacked MHCII and expressed GFP, while MHCII expression on AMs was preserved (**Fig. 4A**). Although 30% of lung DCs appeared to derive from monocytes (**Fig. 1D**), the observation that MHCII was absent on all lung DCs in MHCII^ΔCCR2^ mice is consistent with earlier observations that cDCs are recruited from the bone marrow to the lungs in a CCR2-dependent manner during infection [31, 32].

**Figure 4.**
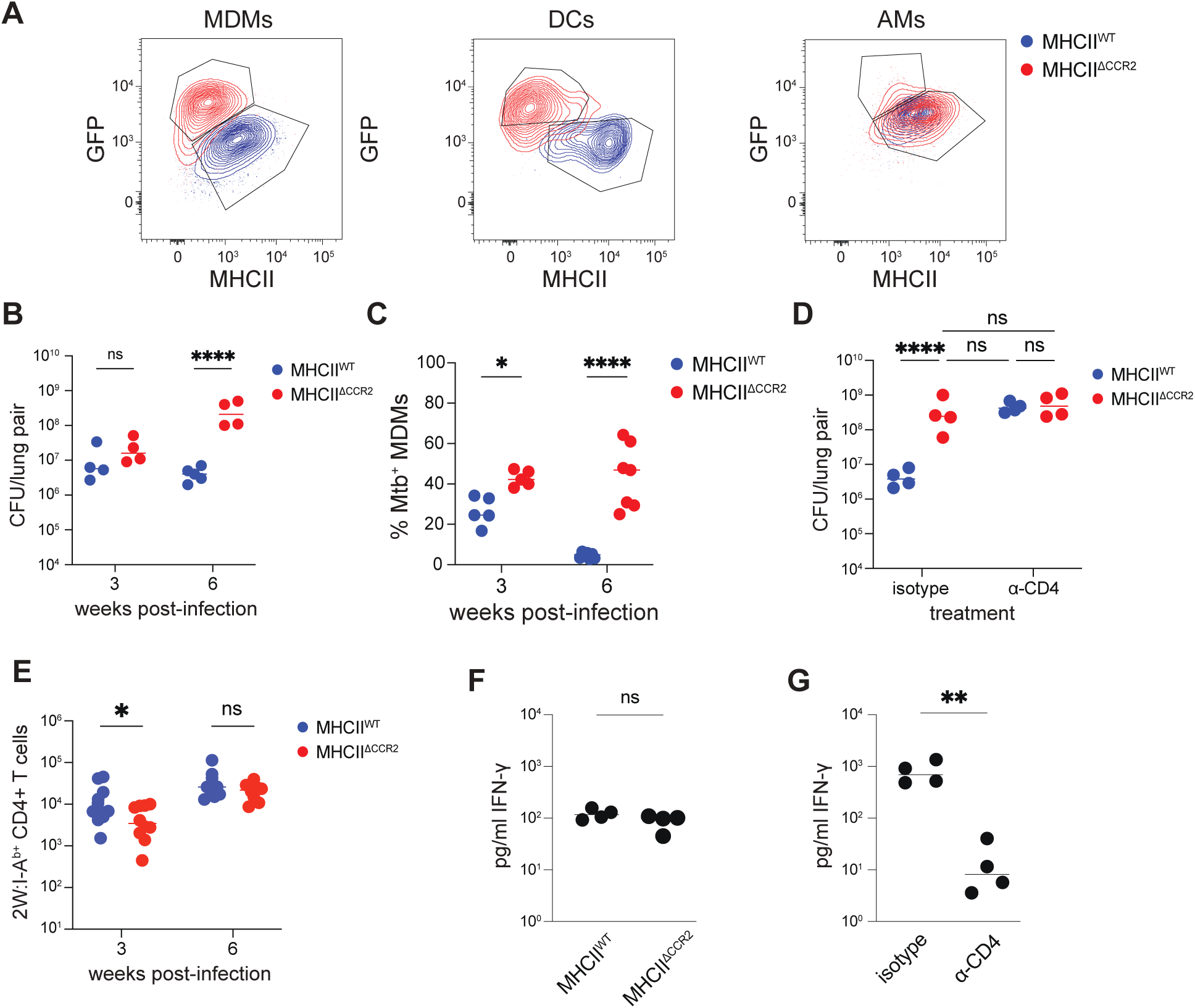
CD4^+^ T cell control of *Mtb* requires MHCII expression by CCR2^+^ cells. (A) GFP and MHCII expression among indicated lung populations from tamoxifen-fed Ccr2-Cre^ERT2^-GFP I-A^b-flox^ mice (MHCII^ΔCCR2^) and Cre^—^ I-A^b-flox^ littermates (MHCII^WT^) at 3 weeks post-infection. (B) CFUs and (C) frequency of infected MDMs in the lungs. (D) CFUs in the lungs of MHCII^WT^ and MHCII^ΔCCR2^ mice treated with α-CD4 or isotype control antibody and infected with *Mtb* for 6 weeks. (E) 2W-specific (2W:I-A^b^ tetramer positive) CD4^+^ T cells in the lungs. (F) IFN-γ abundance in the lungs of MHCII^WT^ and MHCII^ΔCCR2^ mice infected with *Mtb* for 3 weeks. (G) IFN-γ abundance in the lungs of WT mice treated with α-CD4 or isotype control antibodies and infected with *Mtb* for 3 weeks.

MHCII^WT^ and MHCII^ΔCCR2^ mice were then treated with TAM and infected with fluorescent *Mtb* to determine whether antigen presentation by MDMs is needed for CD4^+^ T cells to restrict growth of bacteria within the MDMs. At 3 weeks post-infection, MHCII^ΔCCR2^ mice had slightly more, and at 6 weeks, 100 times more *Mtb* lung CFUs than MHCII^WT^ mice (**Fig. 4B**). In addition, 60% of the MDMs of MHCII^ΔCCR2^ origin in TAM treated chimeric mice contained intracellular *Mtb* at 6 weeks post-infection, whereas very few MDMs of MHCII^WT^were infected (**Fig. 4C**). Thus, overall control of *Mtb* growth in the lungs required MHCII expression by cells with a history of CCR2 expression. The increase in *Mtb* growth observed in MHCII^ΔCCR2^ mice was likely caused by an inability of CD4^+^ T cells to respond to *Mtb* peptide:MHCII complexes on cells with a history of CCR2 expression. This possibility was tested by lung *Mtb* growth in MHCII^WT^ and MHCII^ΔCCR2^ mice treated with either CD4^+^ T cell-depleting antibody or isotype control antibody. At 6 weeks post-infection, isotype control antibody-treated MHCII^ΔCCR2^ mice again contained around 100-fold more *Mtb* CFUs in the lungs compared to isotype-treated MHCII^WT^ mice. By contrast, there was no difference in the *Mtb* lung CFUs between MHCII^WT^ and MHCII^ΔCCR2^ mice treated with CD4^+^ T cell-depleting antibody and both groups contained the same high number of bacteria as isotype-treated MHCII^ΔCCR2^ mice (**Fig. 4D**). Therefore, mice lacking MHCII in CCR2-expressing cells could not generate CD4^+^ T cell-mediated protection against *Mtb* growth.

### *Mtb*-specific Th1 cells develop normally in MHCII^ΔCCR2^ mice despite their inability to restrict *Mtb*

It was possible that the failure of MHCII^ΔCCR2^ mice to restrict *Mtb* growth was due to a defect in the activation, differentiation, or expansion of *Mtb*-specific CD4^+^ T cells. This possibility was tested by tracking CD4^+^ T cells specific for a model peptide antigen called 2W [33] in mice infected with an *Mtb* strain engineered to express this peptide under the control of the ESAT6 promoter. While MHCII^ΔCCR2^ mice contained slightly fewer 2W:I-A^b^-specific CD4^+^ T cells in the lungs compared to MHCII^WT^ mice at 3 weeks post-infection, this difference was no longer observed by 6 weeks post-infection (**Fig. 4E**). Thus, MHCII^ΔCCR2^ mice were not markedly defective in the activation of *Mtb*-specific CD4^+^ T cells or their accumulation in the lungs.

The differentiation state of the *Mtb*-specific CD4^+^ T cells in MHCII^ΔCCR2^ mice was assessed by measuring the Th1 markers CXCR6, Ly-6C, and CX3CR1 [34–37]. No Th1 subset was absent among the 2W:I-A^b^-specific CD4^+^ T cells in the lungs of MHCII^ΔCCR2^ mice compared to MHCII^WT^ mice, although a modest decrease in the number of CXCR6^+^ Ly-6C^+^ cells was observed (**Fig. S4)**. Therefore, MHCII^ΔCCR2^ mice did not have an obvious defect in differentiation of *Mtb*-specific CD4^+^ T cells that could explain the complete lack of CD4^+^ T cell-mediated immunity in MHCII^ΔCCR2^ mice.

Th1 cells secrete IFN-γ upon TCR binding to cognate peptide:MHCII complexes [38]. Because MHCII^ΔCCR2^ mice fail to undergo CD4^+^ T cell-mediated control of *Mtb* growth despite a normal number of *Mtb*-specific Th1 cells in the lungs, it was possible that the Th1 cells failed to secrete IFN-γ because their TCRs were not stimulated by peptide:MHCII complexes on the surface of CCR2^+^. To account for this possibility, IFN-γ abundance was assayed in the lungs of MHCII^WT^ and MHCII^ΔCCR2^ mice at 3 weeks post-infection. As a positive control, IFN-γ was also measured in the lungs of WT mice treated with a-CD4 or isotype control antibody. IFN-γ levels were not significantly different between MHCII^WT^ and MHCII^ΔCCR2^ mice (**Fig. 4F**). By contrast, WT mice treated with a-CD4 had significantly less IFN-γ than isotype control-treated mice (**Fig. 4G**). Thus, the failure of MHCII^ΔCCR2^ mice to restrict *Mtb* growth could not be explained by a global defect in IFN-γ production.

### CMs, NCMs and DCs, but not MDMs, require cognate interactions for optimal receipt of IFN-γ

A single cell RNA sequencing experiment was then performed on lymphocyte-, neutrophil-, and AM-depleted populations from the lungs of MHCII^WT^ and MHCII^ΔCCR2^ mice 3 weeks post-infection with *Mtb* to assess the antimicrobial effect of antigen presentation by CCR2^+^ cells. This time was chosen because the overall *Mtb* burden was similar in the two groups (**Fig. 4D**). When the identities of the captured cells were assessed using the ImmGen database as a reference [39], most of the cells were preliminarily identified as monocytes, macrophages, or DCs, but small populations of contaminating lymphocytes and PMNs were also present (**Fig. S5A**). These contaminants were excluded and the remaining cells were further clustered by transcriptional state.

Analysis of the cells from both MHCII^WT^ and MHCII^ΔCCR2^ mice produced 4 cell clusters (**Fig. 5A**), which likely represented the CMs, NCMs, MDMs and DCs that were observed by flow cytometry (**Fig. 1A-B**). Cluster 3 uniquely expressed *Dpp4* (**Fig. 5B**), which encodes the CD26 protein found on DCs (**Fig. 1B**) [40]. Cluster 1 was enriched for Treml4, which is abundant on NCMs (**Fig. S1A**), and the cluster 1 cells contained other prototypical NCM transcripts including *Cd36*, *Ace* and *Pparg* (**Fig. 5B**) [41]. Cluster 0 had high expression of *Ly6c2* (Ly-6C) but low expression of *H2-Ab1* (MHCII) (**Fig. 5B**), which matches the phenotype of CMs (**Figs. 1B and 1G**). Cluster 0 also was enriched for *Sell* (L-selectin), commonly identified on CMs [42]. Lastly, cluster 2 was enriched for cells expressing both *Ly6c2* and *H2-Ab1*, a feature of MDMs (**Figs. 1B and S1E**), and expressed the inflammatory macrophage genes *Cxcl9*, *Cxcl10* and *Slamf7* (**Fig. 5B**) [43]. When the cells were grouped by genotype, expression of MHCII was attenuated in all four clusters from MHCII^ΔCCR2^ mice (**Fig. S5B**), consistent with the absence of MHCII protein staining on MDMs and DCs from these animals (**Fig. 4A**).

**Figure 5.**
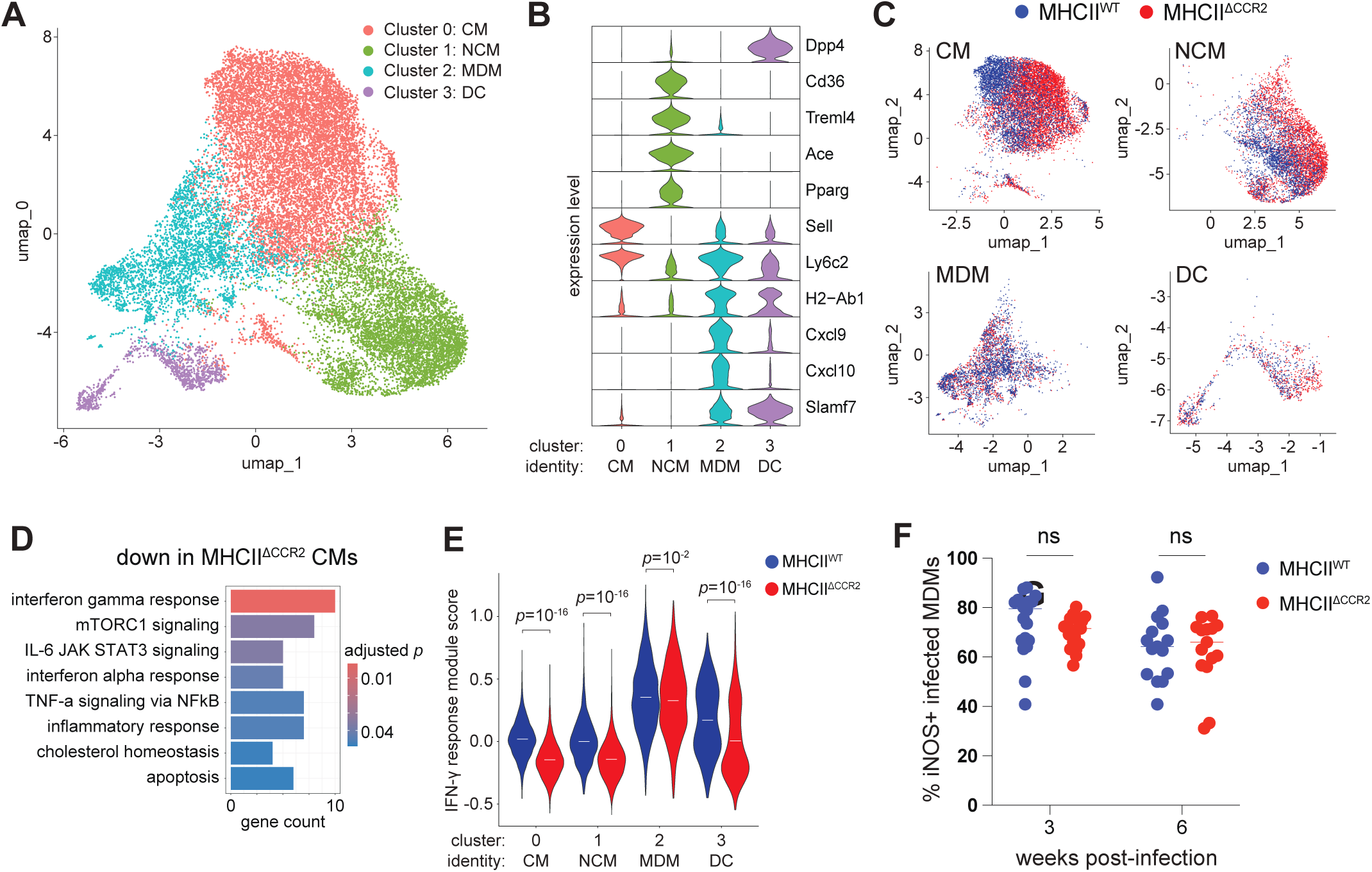
CMs, NCMs and DCs, but not MDMs, require cognate interactions with CD4^+^ T cells for receipt of IFN-γ. (A) Clustering of combined lung myeloid cell populations based on single cell RNA sequencing of lung cells from MHCII^WT^ and MHCII^ΔCCR2^ mice at 3 weeks post-infection. (B) Relative expression of indicated genes (right) among myeloid cell clusters (bottom). (C) UMAP position of CMs, NCMs, MDMs and DCs in MHCII^WT^ compared to MHCII^ΔCCR2^ mice. (D) Biological pathways with decreased expression in CMs from MHCII^ΔCCR2^ compared to MHCII^WT^ mice. (E) IFN-γ-induced gene expression in the indicated clusters from MHCII^WT^ compared to MHCII^ΔCCR2^ mice. (F) Infected cells staining positively for iNOS in indicated mice.

CMs, NCMs, MDMs and DCs were then analyzed for genes that were differentially expressed in MHCII^WT^ and MHCII^ΔCCR2^ mice (**Table S1**). CMs and NCMs originating from the MHCII^ΔCCR2^ mice were shifted in the UMAP relative to CMs and NCMs originating from the MHCII^WT^ mice. By contrast, the position of MDMs and DCs from MHCII^ΔCCR2^ and MHCII^WT^ mice largely overlapped on the UMAP (**Fig. 5C**), indicating that there were stronger MHCII-dependent transcriptional differences in CMs and NCMs compared to MDMs and DCs. Indeed, for genes with an absolute log_2_ fold change in expression of greater than 0.5 and a *p* value of less than 0.05, CMs and NCMs had a greater number of differentially expressed genes between MHCII^WT^ and MHCII^ΔCCR2^ mice (282 and 224) compared to MDMs and DCs (140 and 103) (**Fig. S5C**). This result was unexpected because CMs and NCMs express less MHCII than MDMs and DCs (**Figs. S1E, S5B**).

An attempt was then made to identify biological pathways in CMs, NCMs, MDMs and DCs that were upregulated by cognate interactions with CD4^+^ T cells. Genes with the greatest increase in expression in MHCII^WT^ compared to MHCII^ΔCCR2^ mice (log_2_ fold change of greater than 0.5) were used for an over-representation analysis using the Molecular Signatures Database (MSigDB) Molecular Hallmarks collection as a reference [44] (**Table S2**). Genes expressed by less than 10 percent of cells from MHCII^WT^ sample were excluded to focus on the most prominent differences. For CMs (**Fig. 5D**) and NCMs (**Fig. S5D**), the most over-represented biological pathway was “interferon gamma response”. For DCs, “interferon gamma response” was the third most over-represented pathway (**Fig. S5D**). By contrast, for MDMs, no biological pathways in the Molecular Hallmarks collection were found to be enriched among the differentially expressed genes between MHCII^WT^ and MHCII^ΔCCR2^ mice. Based on these results, it appeared that CMs, NCMs and DCs, but not MDMs, require cognate interactions with CD4^+^ T cells for receipt of IFN-γ. Surprisingly, however, differences in specific biological pathways between the MDMs of MHCII^WT^ and MHCII^ΔCCR2^ mice were not captured by the preliminary analysis.

A more sensitive approach was then used to determine whether MDMs experience any degree of IFN-γ signaling after cognate interactions with CD4^+^ T cells. A module of genes upregulated in response to IFN-γ in the CMs from MHCII^WT^ mice was used to create a IFN-γ response signature, which was applied to all the myeloid cell subsets. MDMs expressed the highest IFN-γ response signature, and its expression was slightly lower in MHCII^ΔCCR2^ mice compared to MHCII^WT^ mice. By contrast, CMs, NCMs and DCs had an overall lower IFN-γ response signature that was strongly attenuated in the cells from MHCII^ΔCCR2^ mice. (**Fig. 5E**).

These results indicated that MDMs do in fact undergo a high degree of IFN-γ-induced gene expression that does not depend on expression of MHCII. This conclusion was supported by the finding that infected MDMs from MHCII^WT^ and MHCII^ΔCCR2^ mice stained similarly with the IFN-γ-induced gene product iNOS (**Fig. 5F**) [45]. Thus, cognate interactions with CD4^+^ T cells enhance IFN-γ signaling in CMs, NCMs, and DCs, but MDMs undergo even greater IFN-γ signaling independently of cognate interactions. Moreover, these results suggest that the failure of MHCII^ΔCCR2^ mice to clear *Mtb* from MDMs is not caused by a defect in the production of iNOS or other IFN-γ-induced gene products in the MDMs.

### A subset of MDMs upregulate glycolysis in response to cognate interactions

A deeper search for pathways other than IFN-γ signaling that could explain the need of MDMs to express MHCII to disinfect their intracellular bacteria was then performed. Although analysis of the myeloid cell subset data did not detect differences between MHCII^+^ and MHCII-MDMs, it was possible that such a difference might be revealed by a narrower comparison of only the *Mtb*-infected MDMs from MHCII^WT^ and MHCII^ΔCCR2^ mice. A new principal component analysis, dimensionality reduction, and clustering of the MDMs was performed separately from the rest of the captured cells. This analysis produced 7 MDM sub-populations (MDM_0 through MDM_6) (**Fig. 6A**). Clusters with cells expressing the iNOS-encoding gene *Nos2* were of special interest as flow cytometry analysis showed that iNOS^+^ MDMs had a much higher fraction of *Mtb* infected cells than iNOS^—^ MDMs (**Fig. 6B**).

**Figure 6.**
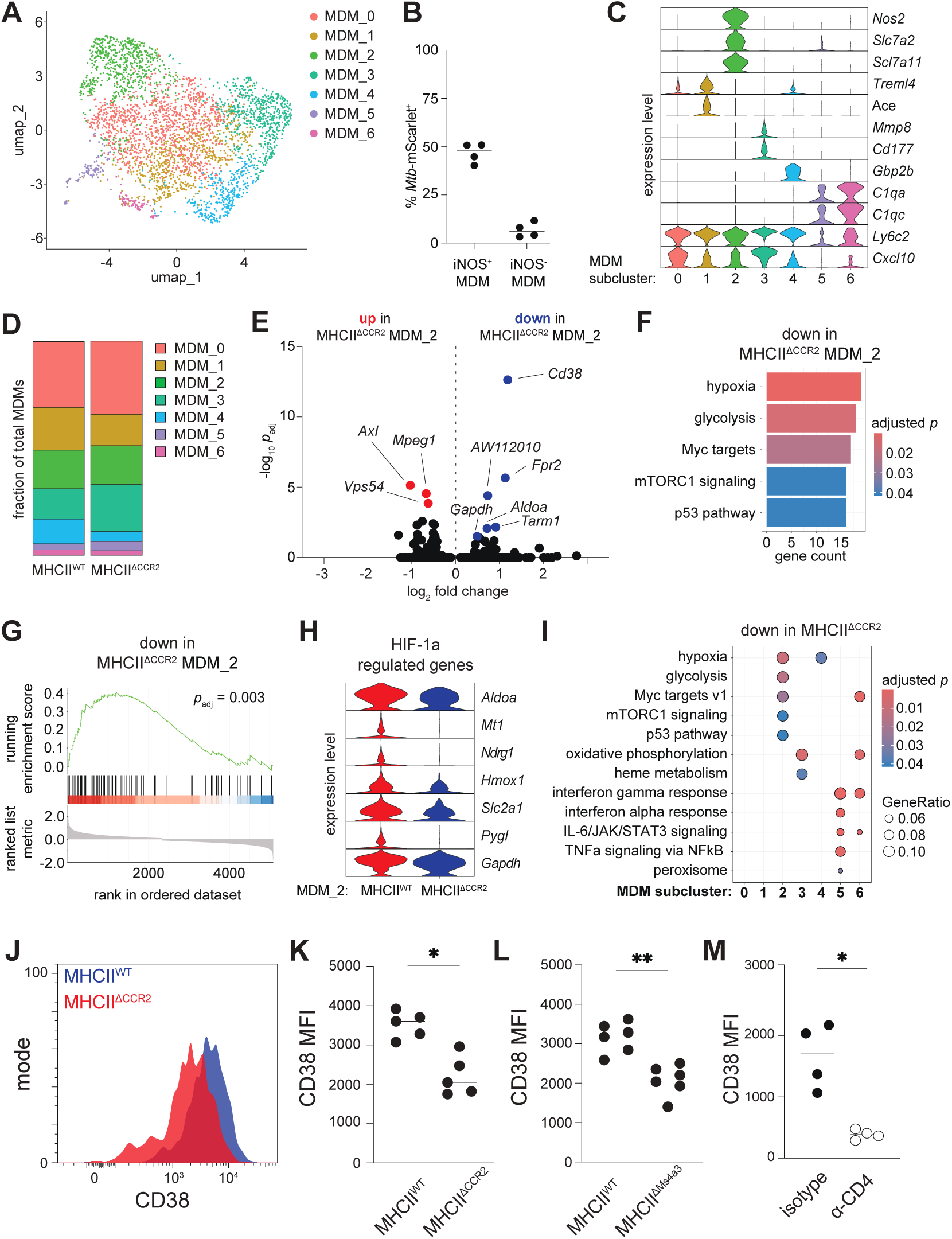
MDMs upregulate glycolysis in response to cognate interactions with CD4^+^ T cells. (A) Subclusters of MDMs among total combined MDMs from MHCII^WT^ and MHCII^ΔCCR2^ mice. (B) Frequency of *Mtb* infection among iNOS^+^ and iNOS-MDMs in WT mice infected for 3 weeks with *Mtb*. (C) Relative expression of indicated genes (right) among MDM subclusters (bottom). (D) Relative frequency of MDM subclusters among total MDMs in MHCII^WT^ and MHCII^ΔCCR2^ mice. (E) Gene expression of MHCII^WT^ MDM_2 cells compared to MHCII^ΔCCR2^ MDM_2 cells. *H2-Ab1* as well as *Rpl9-ps6*, a pseudogene transcript that was absent in all cells from MHCII^ΔCCR2^ samples, were excluded from the plot. (F) Biological pathway expression by MHCII^WT^ MDM_2 cells compared to MHCII^ΔCCR2^ MDM_2 cells. (G) Gene set enrichment analysis (GSEA) of the mSigDB Molecular Hallmarks “glycolysis” geneset applied to MHCII^WT^ MDM_2 cells compared to MHCII^ΔCCR2^ MDM_2 cells. (H) Select HIF-1α-regulated genes with defective expression in MHCII^ΔCCR2^ MDM_2 cells compared to MHCII^WT^ MDM_2 cells. (I) Over-representation analysis of all mSigDB Molecular Hallmarks genesets applied to each subcluster of MDMs from MHCII^WT^ compared to MHCII^ΔCCR2^ mice. (J) CD38 fluorescent antibody staining of infected MDMs in indicated mice. (K-L) Mean fluorescence intensity (MFI) of CD38 staining of infected MDMs in indicated mice. (M) CD38 MFI of infected MDMs in WT mice treated with α-CD4 or isotype control antibody. Data in (J) through (M) was collected at 3 weeks post-infection.

Cells in MDM_2 had a transcriptional signature that was enriched for synthesis of nitric oxide (NO), with abundant transcripts for *Nos2* and the arginine acquisition-related genes *Slc7a2* and *Slc7a11* (**Fig. 6C**) [46]. Cells in MDM_0, which was adjacent to MDM_2 in the UMAP (**Fig. 6B**), were transcriptionally similar to those in MDM_2 except for expression of the NO-promoting genes (**Fig. 6C**). MDM_1 contained cells that had an NCM-like phenotype based on expression of *Treml4* and *Ace*, but, in contrast to NCMs, expressed *Ly6c2* at similar abundance to other MDM populations. Cells in MDM_3 expressed *Mmp8* and *Cd177*, genes associated with activated PMNs [47, 48] and MDM_4 differed from other cells with respect to expression of *Gbp2b*, encoding a GTPase whose family is associated with *Mtb* protection [49]. MDM_5 and MDM_6 contained minor populations that were enriched for C1q transcripts and expressed lower levels of *Ly6c2* and *Cxcl10* than the other MDM subclusters (**Fig. 6C**). Overall, the cells in the MDM subclusters had similar frequencies in MHCII^WT^ and MHCII^ΔCCR2^ mice, although MHCII^ΔCCR2^ mice hadd fewer cells of the *Gbp2b*-expressing subcluster MDM_4 (**Fig. 6D**).

The MDM_2 subcluster, which was enriched for *Nos2* transcripts (**Fig. 6C**) and therefore contained the majority of *Mtb*-infected MDMs (**Fig. 6B**), was analyzed further. *Cd38*, *Aldoa*, *Gapdh, Fpr2*, *Tarm1*, and *Aw112010* were among the most significantly downregulated genes in MDM_2 cells in MHCII^ΔCCR2^ mice, whereas *Axl*, *Mpeg1*, and *Vps54* were significantly upregulated (**Fig. 6E**) (**Table S3**). Over-representation analysis identified genes related to hypoxia and glycolysis as significantly enriched among the genes with defective expression in MHCII^ΔCCR2^ mice. Targets of the transcription factors Myc and p53 were also enriched, as were genes involved in mTORC1 signaling (**Fig. 6F**).

The over-representation analysis results were notable because upregulation of glycolytic metabolism in macrophages is strongly associated with *Mtb* disinfection [18, 50, 51]. Moreover, genes promoting glycolysis are induced by Myc, mTORC1, and the hypoxia-responsive transcription factor HIF-1α [52, 53]. The over-representation analysis was validated using a more sensitive technique of gene set enrichment analysis (GSEA) [44]. Genes positively associated with glycolysis were significantly enriched among the ranked list of downregulated genes in MHCII^ΔCCR2^ MDM_2 cells (**Fig. 6G**). Many of the glycolysis-related genes identified by GSEA are known to be direct targets of HIF-1α, which is essential for immunity to *Mtb* in mice [50]. HIF-1α -regulated genes with impaired expression in MDM_2 cells from MHCII^ΔCCR2^ mice included *Aldoa*, *Mt1*, *Ndrg1*, *Hmox1*, *Slc2a1*, *Pygl*, and *Gapdh* (**Fig. 6H**) [54–57]. MHCII-dependent upregulation of glycolysis was a unique feature of MDM_2 cells because a comparative over-representation analysis showed that MHCII-dependent upregulation of genes associated with hypoxia, glycolysis, Myc, mTORC1 and p53, while observed in MDM_2 cells, was not consistently observed for the other MDM subclusters. Instead, MDM_3 cells from MHCII^ΔCCR2^ mice were defective in the expression of genes related to oxidative phosphorylation and heme metabolism, while the small subclusters MDM_5 and MDM_6 were defective in the expression of peroxisome and inflammatory cytokine-induced genes in MHCII^ΔCCR2^ mice (**Fig. 6I**) (**Table S4**). Together, the results suggest that MDM_2 cells that are most likely infected with *Mtb* upregulate glycolytic metabolism upon cognate interactions with CD4^+^ T cells.

### CD4^+^ T cells and MHCII expression by MDMs are required for upregulation of CD38 in infected MDMs

The most significantly downregulated gene in MHCII-deficient MDM_2 cells was *Cd38* (**Fig. 6E**), an enzyme that promotes glycolysis by stabilizing HIF-1α [58]. CD38 is also expressed on the cell surface and is readily detected by flow cytometry. This feature of CD38 was used to perform a final validation that cognate MHCII-mediated interactions with CD4^+^ T cells induce a glycolytic program within infected MDMs. Indeed, infected MDMs from MHCII^ΔCCR2^ mice had significantly weaker CD38 staining compared to MHCII^WT^ mice (**Fig. 6J-K**). An additional approach was then used to verify MHCII-dependent induction of CD38 in MDMs by generating *Ms4a3^Cre^ H2-Ab1^flox/flox^* (MHCII^ΔMs4a3^) mice, in which cells of the monocyte and granulocyte lineages delete the gene encoding the I-A^b^ beta chain. As with MHCII^ΔCCR2^ mice, CD38 staining was lower on the infected MDMs of MHCIIdMs4a3 mice compared to the MDMs of MHCII^WT^ mice (**Fig. 6L**). Finally, depletion of CD4^+^ T cells in WT mice led to the reduction of CD38 on MDMs (**Fig. 6M**). Together, the results demonstrated that CD4^+^ T cells and MHCII expression by MDMs are both required for upregulation of CD38 in infected MDMs.

## Discussion

The goal of this study was to determine how Th1 cells control *Mtb* infection in the lungs and to clarify the role of IFN-γ in this process. The results point to an “attract and kill” model in which Th1 cells produce IFN-γ in the lungs to attract MDMs, which are preferentially infected by *Mtb*. The infected MDMs subsequently present *Mtb* peptide:MHCII complexes on their surfaces and engage CD4^+^ T cells in cognate interactions. These cognate interactions promote disinfection by inducing glycolytic metabolism in the MDMs. Thus, Th1 cells induce a cycle of MDM recruitment, *Mtb* phagocytosis and MDM disinfection that results in bacterial control.

It was important to accurately identify the myeloid cells that were in the lungs of *Mtb*-infected mice and contained intracellular bacteria to achieve the goal of this study. Earlier work identified a mixture of poorly characterized myeloid populations and IMs as the primary population of *Mtb*-infected myeloid cells within the lungs of *Mtb*-infected mice. The nature of many of these populations cells was ambiguous, however, because of co-expression of macrophage and DC surface markers. By staining with CX3CR1 and Ly-6C antibodies, we observed that NCMs constitute a significant fraction of cells within *Mtb*-infected lungs. NCMs likely account for the CD11c^int^ MHCII^int^ cells that have been consistently observed but whose identity and ontogeny were unknown [14–18]. We found, however, that NCMs and CMs were present in the blood component of the lung tissue and thus unlikely to play a direct role in restricting *Mtb* growth in the lung interstitium where *Mtb* lesions are located. Instead, we found that CX3CR1^lo^ CD11c^hi^ Ly-6C^hi^ CD26^lo^ CM-derived macrophages, which were distinguished from CX3CR1^lo^ CD11c^hi^ Ly-6C^lo^ CD26^hi^ DCs, had the features previously ascribed to IMs and were the main subset of *Mtb*-infected mononuclear phagocytes cells in the lung interstitium.

The fact that recruitment of CM-derived MDMs into the lung tissue requires CD4^+^ T cell-derived IFN-γ, but not expression of the IFN-γ receptor, indicates that IFN-γ promotes MDM recruitment by an indirect mechanism. In other models, direct IFN-γ signaling to endothelial cells was required for vascular permeabilization and infiltration of leukocytes into inflamed tissue [59]. IFN-γ also induces the expression of chemokines and endothelial surface proteins that promote extravasation [60, 61]. Thus, IFN-γ secreted by *Mtb-*specific CD4^+^ T cells may increase chemokine expression or activate lung endothelial cells to promote CM migration from the blood into the lung tissue. In any case, the recruitment of MDMs by CD4^+^ T cells may be an evolved mechanism to lure *Mtb* bacteria into the host cell most capable of engaging CD4^+^ T cells and receiving microbicidal signals.

While prior studies showed that antigen presentation by MDMs is dispensable for the initial activation of *Mtb*-specific CD4^+^ T cells in the lung-draining lymph node [30], it was possible that antigen presentation by MDMs maintained *Mtb-*specific CD4^+^ T cell survival or IFN-γ production in the lungs. This possibility was excluded, however, by the observation that the lungs of mice lacking the MHCII gene in all CCR2^+^ cells had normal amounts of *Mtb*-specific CD4^+^ T cells and IFN-γ. Thus, cells other than CMs, NCMs, MDMs or CCR2-expressing DCs present antigens to *Mtb*-specific CD4^+^ T cells within the lungs to maintain CD4^+^ T cell survival and IFN-γ secretion. Alternatively, cytokines such as IL-12 or IL-18 produced within *Mtb*-infected lungs could maintain CD4^+^ T cell survival and IFN-γ secretion independently of TCR signaling, as has been reported elsewhere [62].

Although cognate interactions between MDMs and CD4^+^ T cells were dispensable for maintaining the CD4^+^ T cells, they were required for the ability of the CD4^+^ T cells to induce glycolytic gene expression in the MDMs. The metabolic shift of macrophages towards glycolysis, mediated in large part by HIF-1α, is required for immunity to *Mtb* and other intracellular bacterial pathogens [63, 64]. The mechanisms by which glycolysis in macrophages supports antimicrobial immunity are poorly understood but may include limiting metabolites that support bacterial growth, producing microbicidal metabolites, or meeting energy demands for additional biochemical processes in the macrophage. Notably, CD4^+^ T cells extracted from *Mtb-*infected mice were shown to suppress *Mtb* growth within bone marrow-derived macrophages in a HIF-1α-dependent manner and induced a similar set of hypoxia- and glycolysis-related genes [65]. Our study supports these earlier observations and further establishes that induction of glycolysis in infected MDMs is mediated by cognate interactions with CD4^+^ T cells *in vivo*. Interactions between ligands that are acutely induced on CD4^+^ T cells by TCR recognition of peptide:MHCII complexes and receptors on MDMs that display such complexes are good candidates for the transducers of the signals that induce glycolysis in MDMs.

## Acknowledgements

The authors acknowledge the following funding sources: R01AI103760 (MKJ), 2T32HL007062 (SHB), R01AI173780 (TDB), and K08AI150425 (TDB). The University of Minnesota Flow Cytometry Resource Core and Biosafety Level 3 Program were critical to the study. The Roy D. Carver Biotechnology Center at the University of Illinois Urbana-Champaign (Alvaro Gonzalo Hernandez, Chris Wright) sequenced the libraries for single-cell RNA sequencing. The authors also thank Thamotharampillai Dileepan for providing tetramer reagents, Jesse Williams for criticism related to the study and the manuscript, and Gregory Vercellotti for overseeing the T32 training program.

## Author Contributions

Conceptualization, S.H.B., T.D.B., and M.K.J.; Methodology, S.H.B. and M.K.J.; Formal Analysis, S.H.B.; Investigation, S.H.B. and C.E.R.; Data Curation, S.H.B.; Writing–Original Draft, S.H.B.; Writing–Review & Editing, S.H.B., T.D.B., and M.K.J.; Visualization, S.H.B.; Supervision, M.K.J.; Funding Acquisition, M.K.J.

## Declaration of interests

The authors declare no competing interests.

## Supplemental information

Table S1. Excel file containing additional data too large to fit in a PDF, related to Figures 5C and S5C.

Table S2. Excel file containing additional data too large to fit in a PDF, related to Figures 5D and S5D.

Table S3. Excel file containing additional data too large to fit in a PDF, related to Figure 6E.

Table S4. Excel file containing additional data too large to fit in a PDF, related to Figure 6I.

## Materials and methods

### Animals

Mice were housed under specific pathogen-free conditions in accordance with University of Minnesota Institutional Animal Care and Use Committee guidelines. C57BL/6J, and B6.SJL-Ptprca Pepcb/BoyJ (CD45.1) mice were bred in-house. The following mouse lines were purchased from Jackson Laboratories and bred in-house: B6.129S2-Tcratm1Mom/J (*Tcra^-/-^*), C57BL/6J-Ms4a3em2(cre)Fgnx/J (*Ms4a3*^Cre^), B6.Cg-Gt(ROSA)26Sortm9(CAG-tdTomato)Hze/J (tdTomato^LSL^), B6.129S7-Ifngtm1Ts/J (*IFNg*^-*/-*^), B6.129S7-Ifngr1tm1Agt/J (*IFNgR1^-/-^*), C57BL/6-Ccr2em1(icre/ERT2)Peng/J (*Ccr2-CreERT2-GFP*), B6.129X1-H2-Ab1b-tm1Koni/J (*H2-Ab1^fl/fl^*). *Ms4a3*^Cre^ and tdTomato^LSL^ mice were bred for one generation to produce mice heterozygous for both alleles. MHCII^WT^ and MHCII^ΔCCR2^ mice were generated by crossing *Ccr2-CreERT2-GFP* and *H2-Ab1^fl/fl^* for several generations to produce litters of *H2-Ab1^fl/fl^* mice that were either heterozygous or null for the *Ccr2-CreERT2-GFP* allele. Littermates were used for experiments. The same approach was used to generate MHCII^dMs4a3^ mice and MHCII^WT^ littermate controls. Mice used for experiments were 4-8 weeks old with the exception of bone marrow chimeric mice, which were 4-8 weeks old at the time of irradiation and 14-18 weeks old at the time of infection. MHCII^WT^ and MHCII^ΔCCR2^ mice were fed with chow containing tamoxifen (500 mg/kg diet formulation, Inotiv Teklad) starting on the day of *Mtb* challenge. Mice were age-matched for experiments and approximately equal proportions of male and female mice were used.

### Bacterial strains and plasmids

The *Mycobacterium tuberculosis* strain H37Rv was transformed with a chromosomally integrating plasmid expressing a variant of mScarlet fluorescent protein [66] to generate strain *Mtb*-mScarlet. This plasmid is selectable by hygromycin resistance, using the mycobacterial optimized promoter (MOP) to drive mScarlet expression and a Giles phage integration module [67].

To make fluorescent H37Rv expressing 2W peptide, an open reading frame encoding H37Rv EsxA (Rv3875) with a C-terminal 4 x glycine linker, followed by 2W peptide [33], followed by a FLAG peptide was inserted into pMN402 [68]. pMN402 contains the promoter region of H37Rv Hsp60 (Rv0440) upstream of the insertion site. The promoter and coding sequence were subcloned into pTT1B, which encodes gentamycin resistance and chromosomally integrates into the *Mtb* L5 integration locus [69]. The resulting plasmid (pTT1B-EsxA-2W) was transformed into *Mtb*-mScarlet by electroporation and transformants were selected on hygromycin- and gentamycin-containing agar plates.

### Mouse infections

Mice were infected with a low dose (100-200 CFU/lung pair) of *Mtb* via aerosol exposure. *Mtb* strains were grown from frozen stocks in 7H9 broth supplemented with bovine serum albumin, dextrose and NaCl. For preparing the bacterial inoculum, *Mtb* was grown to an optical density at 600 nm (OD600) of 0.4-0.6, washed in PBS-T (PBS containing 0.5% Tween-80) and diluted to an OD600 of 0.005. The inoculum was nebulized for aerosol delivery using the Glas-Col inhalation exposure system. The infectious dose was monitored by plating total lung homogenate the day after aerosol exposure. For CD4^+^ T cell depletion, α-CD4 or isotype control antibodies were intraperitoneally injected weekly starting on the day of infection. For CSF1R blockade, α-CD115 or isotype control antibodies were intraperitoneally injected on days 14, 16, 18, and 20.

### Processing of infected lungs for flow cytometry and cell sorting

Mice were euthanized using carbon dioxide inhalation in accordance with University of Minnesota Institutional Animal Care and Use Committee guidelines. Mouse lungs were placed in IMDM complete media (IMDM supplemented with GlutaMAX, pyruvate, non-essential amino acids, 10% fetal bovine serum and 200 µM beta-mercaptoethanol) (Gibco) and homogenized using a GentleMACS tissue dissociator (Miltenyi). Homogenates were strained through a 70 µm filter to generate single cell suspensions.

### Cell staining and counting for flow cytometry and cell sorting

Single cell suspensions were split into multiple volumes for T cell and myeloid cell staining. For T cell staining, cells were stained with flow cytometry antibodies (1:50 dilution), fluorophore-conjugated peptide:MHCII tetramers (1:100 dilution) and viability dye (1:500 dilution) for 20 minutes at room temperature. For myeloid cell staining, cells were stained with flow cytometry antibodies (1:50 dilution) and viability dye (1:500 dilution) for 20 minutes at room temperature. For intracellular staining with α-iNOS, cells were treated with fixation/permeabilization solution (BD), washed with permeabilization wash buffer (Tonbo), and stained in permeabilization wash buffer for 20 minutes at room temperature. For all samples subject to conventional flow cytometry analysis, cells were fixed overnight in 5% formalin solution at 4°C for sterilization. For i.v. labeling experiments, mice were injected retro-orbitally with 2.5 µg of fluorophore-conjugated α-CD45 3 minutes prior to euthanasia. To obtain cell counts, a master mix of AccuCheck Counting Beads (ThermoFisher Scientific) in flow cytometry buffer was applied in equal volumes to the fixed cell samples immediately prior to analysis on the flow cytometer. For each experiment, including time course experiments, the same counting bead master mix was used for samples and time points in the experiment.

### Monocyte and CD4^+^ T cell purification and adoptive transfer

Monocytes were purified from donor bone marrow using the Stemcell Mouse Monocyte Isolation Kit according to the manufacturer’s guidelines and resuspended in PBS. Monocytes purified from donor mice were retro-orbitally injected into *Mtb-*infected recipient mice at a 1:1 ratio (2-4 x 10^6^ monocytes per recipient mouse). For purification and adoptive transfer of CD4^+^ T cells, lymph nodes and spleens of infection-naïve donor mice were pooled, homogenized using a GentleMACS tissue dissociator and strained through a 70 µm filter to generate a single-cell suspension. CD4^+^ T cells were purified from the spleen and lymph node suspensions by negative selection and magnetic separation as follows. Cells in IMDM complete media were incubated with biotinylated antibodies against CD11b, CD11c, CD16/32, B220, CD8a, Ter119, NK1.1 and F4/80 (1:100 dilution) for 15 minutes at room temperature. Stained cells were mixed with streptavidin-conjugated magnetic beads (Stemcell) for 3 minutes, after which a larger volume of complete media was added and the samples were applied to a magnet (Stemcell) for 3 minutes. CD4^+^ T cells were collected by recovering the unbound fraction. CD4^+^ T cells purified from donor mice were resuspended in PBS and retro-orbitally injected into infection-naïve recipient mice at a 1:1 ratio (approximately 20 million CD4^+^ T cells per recipient mouse). The CD4^+^ T cell recipient mice were rested for 7 days before challenging with *Mtb*.

### Enumeration of bacterial colony-forming units

Mouse lungs were homogenized as described above and and an aliquot of homogenate was mixed 1:1 with PBS-T without prior straining. Serial dilutions were prepared in PBS-T and plated onto 7H11 agar containing Middlebrook OADC supplement.

### Cytokine and chemokine quantification

*Mtb*-infected mouse lungs were homogenized as described above. A fraction of the homogenate was centrifuged at 10,000 x g for 5 minutes at 4°C. The supernatant fraction was transferred to a Spin-X 22 µm filter centrifuge tube (Corning) and centrifuged for 30 minutes at 4°C. The filtrate was collected and stored at −20°C until assayed. Cytokines and chemokines were measured in the samples using the LEGENDplex Mouse Inflammation Panel and LEGENDplex Mouse Proinflammatory Chemokine Panel bead-based ELISA kits (BioLegend). Concentrations were obtained using standards provided in the kits according to the manufacturer guidelines.

### Generation of mixed bone marrow chimeric mice

Using an X-ray irradiator, mice were irradiated with 5 x Gly, rested for 24 hours, and irradiated a second time with 5 x Gly. Bone marrow was collected from donor mice in DMEM media (Gibco) containing 10% fetal bovine serum and counted using a hemocytometer. 2.5 x 10^5^ bone marrow cells from each donor were pooled and transferred into irradiated recipient mice via retro-orbital injection. Mice were reconstituted for approximately 10 weeks prior to infection.

### Single cell RNA sequencing

To obtained sorted lung cells enriched for monocytes, MDMs and DCs, single cell suspensions were generated from the lungs of 2 MHCII^WT^ mice (1 male and 1 female) and 2 MHCII^ΔCCR2^ mice (1 male and 1 female) and kept at 4°C in IMDM complete media. Cells were stained with antibodies against Thy1.2, B220, CD19, Ly-6G, Siglec-F, CD11b, and NK1.1 (1:25 dilution) and viability dye (1:500 dilution) for 30 minutes.

Cells staining positively for CD11b and negatively for the remaining markers were sorted using a Sony MA-900 cell sorter. Cells were sorted into IMDM complete media with 40% FBS. Approximately 30,000 cells per sample were recovered from the sorter. Sorted cell samples were processed to obtain RNA in separate wells of the same microfluidics chip using the Chromium Next GEM Single Cell 3’ Gene Expression kit and Chromium X controller (10X Genomics). cDNAs were amplified and libraries barcoded by sample ID were generated according to the manufacturer instructions. Libraries were sequenced by the DNA Services lab at the University of Illinois at Urbana-Champaign. For sequencing, the libraries were pooled, quantitated by qPCR and sequenced on 2 10B lanes with 28 x 150 nucleotide reads on a NovaSeq X Plus with V1.0 sequencing kits (Illumina). The samples each yielded 350-450 million paired reads.

### Analysis of RNA sequencing data

Analysis of single cell RNA sequencing data was performed in R. Using the Seurat package [70], mitochondrial gene features were identified and cells with a mitochondrial feature content of greater than 5% were counted as non-viable and removed from the analysis. Cells with outlying feature counts (under 500 or over 4000 features) were also removed. The feature counts for each cell were normalized and scaled using default Seurat parameters as follows. For normalization, the count of each feature was divided by total counts for that cell and multiplied by 10,000. For scaling, features were centered to have a mean of 0 and scaled by the standard deviation of that feature. Scaled data was used to identify variable features, run principal component analysis, and perform differential gene expression analysis. SingleR [71] was used to identify contaminating lymphocytes, NK cells and PMNs by integrating reference data from ImmGen [39]. The clusterProfiler package [72] was used to perform over-representation analysis and GSEA. For GSEA, differentially expressed genes were ranked in decreasing order of log_2_ fold change. The IFN-γ response signature (**Fig. 5E**) consisted of genes within the mSigDB “interferon gamma response” geneset [44] that were enriched in CMs from MHCII^WT^ versus MHCII^ΔCCR2^ mice based on over-representation analysis.

### Statistical methods

Statistical comparison of 2 groups was performed using a parametric 2-tailed t test. For comparison of more than 2 groups, ANOVA was used with correction for multiple comparisons.

## Supplemental information

**Table S1.**Excel file containing additional data too large to fit in a PDF, related to Table S2. Excel file

---

**Figure S1.**
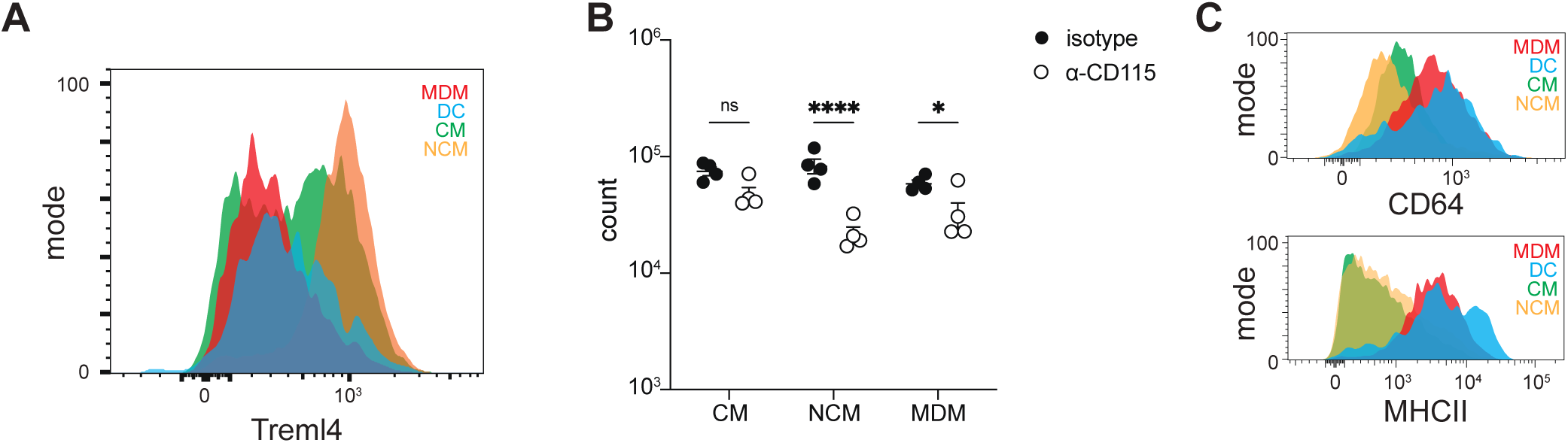
(A) Treml4 antibody staining of indicated cells in the lungs of infected WT mice. (B) Number of indicated cells in the lungs of mice treated for 1 week with CSF1R blocking antibody (α-CD115) or isotype control antibody. (C) CD64 and MHCII antibody staining of indicated cells in the lungs of infected WT mice. Data was collected at 3 weeks post-infection.

**Figure S2.**
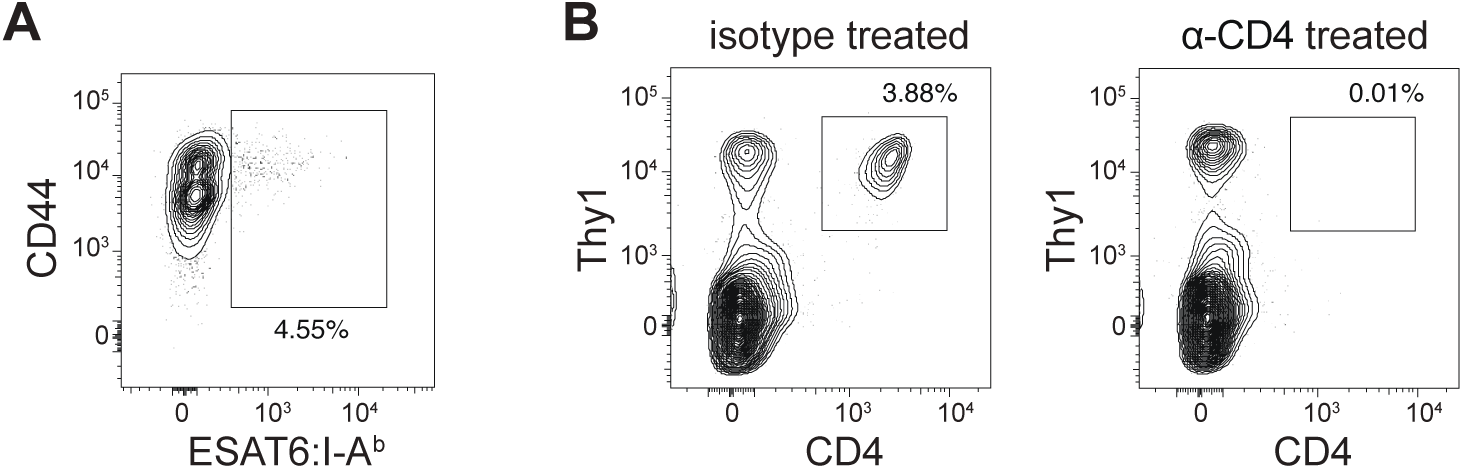
(A) Staining of lung CD4^+^ T cells with ESAT6:I-A^b^ tetramer. (B) Lung CD4^+^ T cell abundance in mice treated with α-CD4 or isotype control antibody. Data was collected at 3 weeks post-infection.

**Figure S3.**
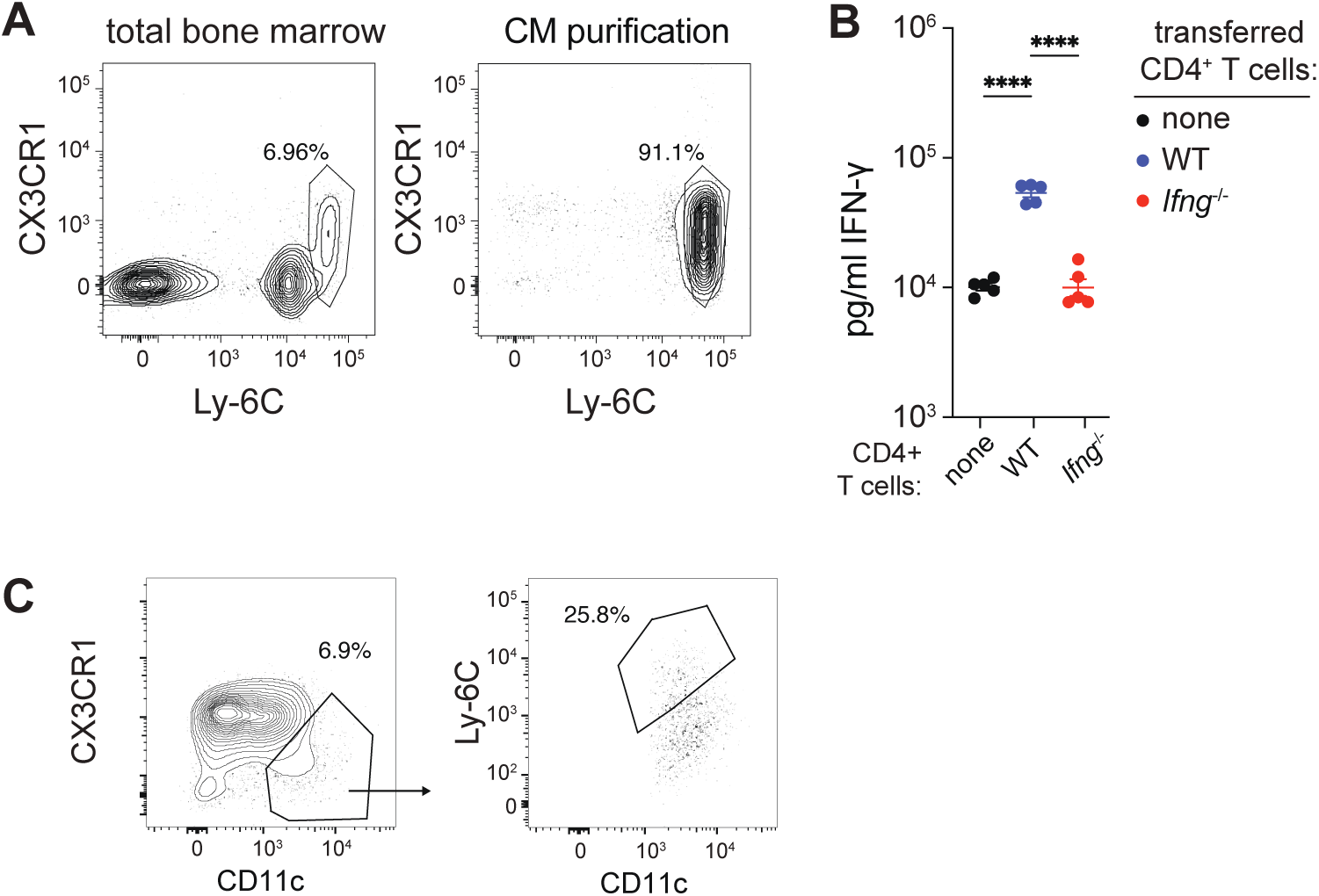
(A) Bone marrow CMs before (left) and after (right) purification. (B) Abundance of IFN-γ in the lungs of T cell-deficient mice given no CD4^+^ T cells, WT CD4^+^ T cells, or IFN-γ-deficient (*Ifng*^—/—^) CD4^+^ T cells and infected for 4 weeks with *Mtb*. (C) Flow cytometry gating to identify MDMs in the lungs of mixed bone marrow chimeric mice.

**Figure S4.**
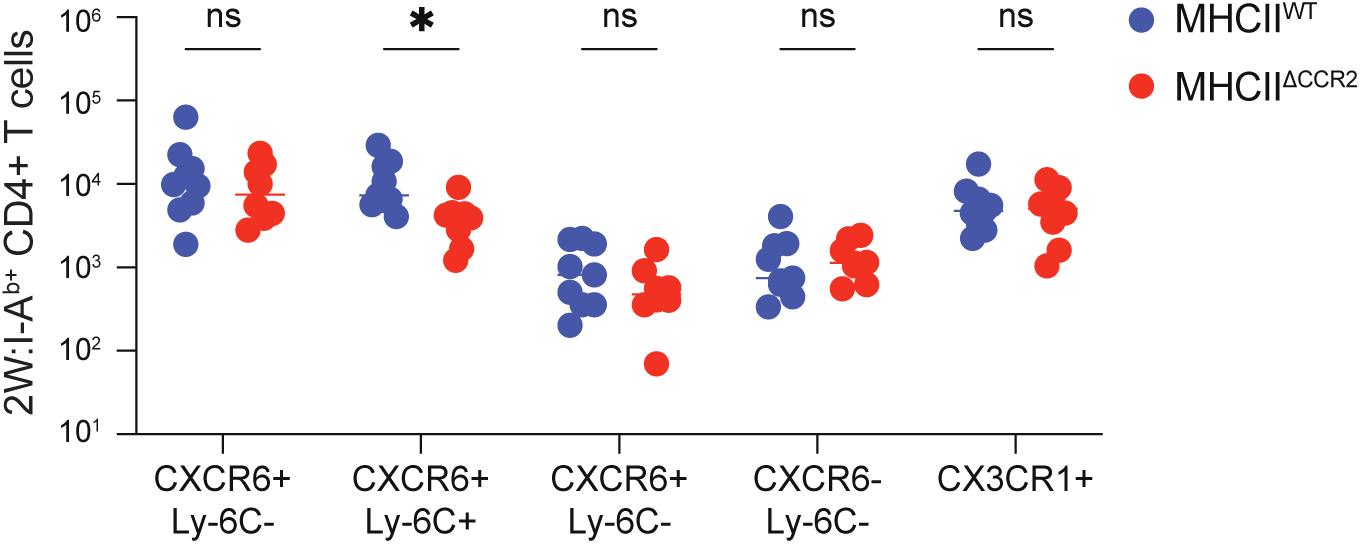
Abundance of subsets of 2W:I-A^b^ tetramer positive CD4^+^ T cells in the lungs of indicated mice at 3 weeks post-infection.

**Figure S5.**
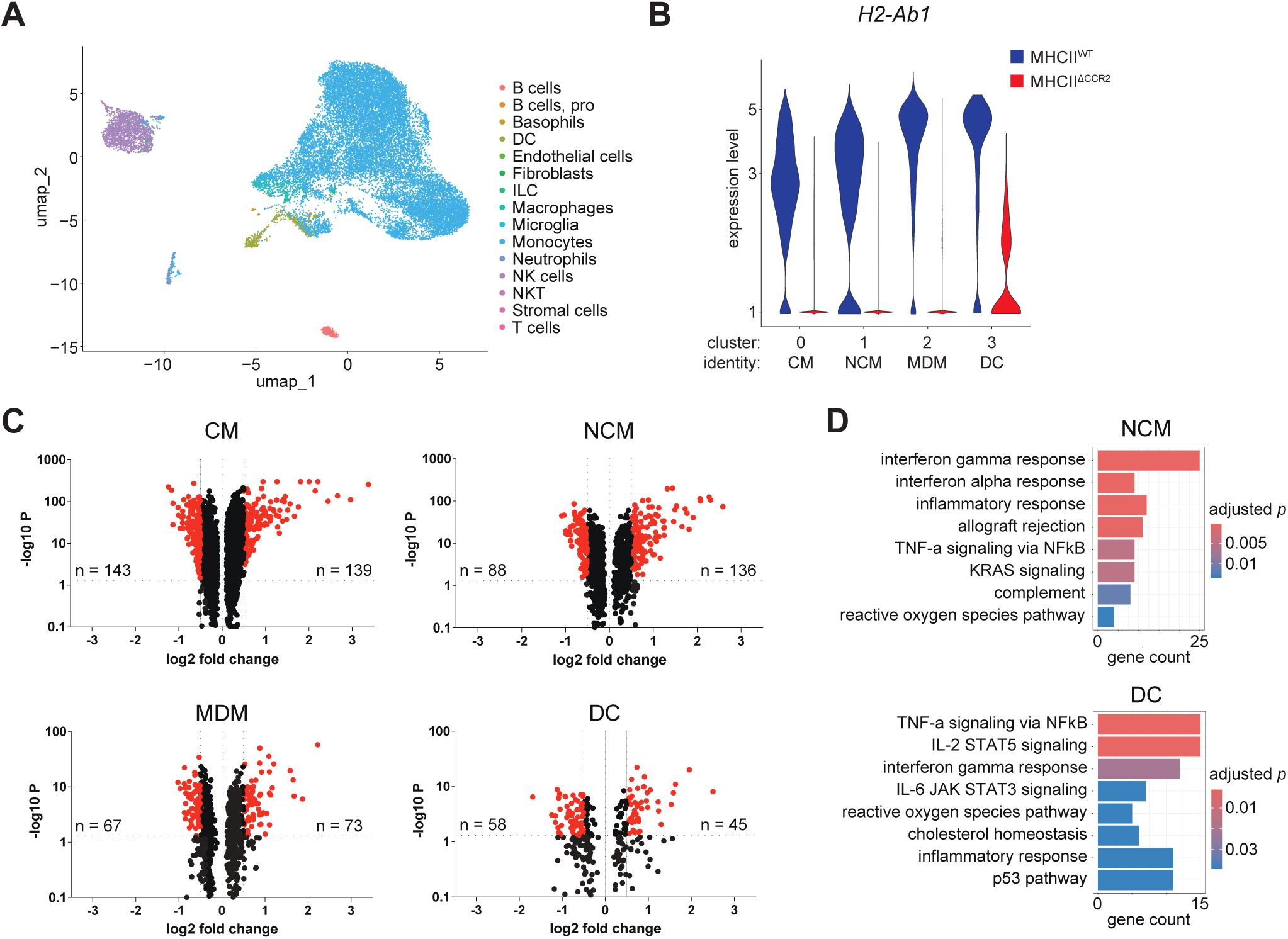
ImmGen annotation of total sorted lung cells after single cell RNA sequencing. (B) *H2-Ab1* expression by indicated cell clusters in MHCII^WT^ compared to MHCII^ΔCCR2^ mice. (C) Gene expression by indicated clusters from MHCII^WT^ (positive log_2_ fold change) compared to MHCII^ΔCCR2^ (negative log_2_ fold change) mice. (D) Biological pathway expression by NCMs (top) and DCs (bottom) from MHCII^WT^ mice compared to MHCII^ΔCCR2^ mice.

## Notes

### Competing Interest Statement

The authors have declared no competing interest.

